# Variation in drug penetration does not account for the natural resistance of *Mycobacterium abscessus* biofilms to antibiotic

**DOI:** 10.1101/2024.04.16.589735

**Authors:** Winifred C. Akwani, Paulina Rakowska, Ian Gilmore, Mark Chambers, Greg McMahon, Suzie Hingley-Wilson

## Abstract

*Mycobacterium abscessus*, an inherently drug-resistant, opportunistic, nontuberculous mycobacterium (NTM) predominantly causes pulmonary infections in immunocompromised patients, notably those with cystic fibrosis. *M. abscessus* subspecies display distinct colony morphologies (rough and smooth), with the prevalent view that *M. abscessus* (smooth) is a persistent, biofilm-forming phenotype, whilst *M. abscessus* (rough) is unable to form biofilms. Biofilm formation contributes to persistent infections and exhibits increased antibiotic resistance.

We used the chemical mapping technique, nanoscale secondary ion spectrometry (NanoSIMS), to investigate if variations in the biofilm morphology and antibiotic penetration account for the antibiotic susceptibility amongst *M. abscessus* subspecies, contributing to increased antimicrobial resistance (AMR) and potentially explaining the protracted treatment duration.

The susceptibility to bedaquiline (BDQ) of *M. abscessus* grown as planktonic bacilli and biofilms was measured. The minimum biofilm eradication concentration (MBEC) of BDQ was 8-16 times higher (2-4µg/ml) compared with the minimum inhibitory concentration (MIC) (0.25µg/ml), indicating reduced efficacy against biofilms.

Correlative imaging with electron microscopy revealed that *M. abscessus* (irrespective of the colony morphotype) formed biofilms and that BDQ treatment influenced biofilm morphology. We determined that *M. abscessus* morphotypes exhibit differential uptake of the antibiotic BDQ in biofilms. *M. abscessus* subsp. *abscessus* (smooth) biofilms exhibited the least uptake of BDQ, whereas *M. abscessus* subsp. *bolletii* biofilms showed the greatest antibiotic penetration.

NanoSIMS analysis revealed no correlation between antibiotic penetration and drug efficacy within the biofilm. This challenges the previous assumption linking biofilm architecture to drug efficacy. Investigating other biofilm characteristics like antibiotic persistence could lead to enhanced treatment approaches.

**Significance Statement:** *Mycobacterium abscessus* is an increasingly prevalent pathogen, most often causing lung infections in immunocompromised individuals. Their distinct morphotypes and biofilm-forming capabilities contribute to persistent infections, rendering them challenging to treat with increased antibiotic resistance. This research demonstrates that the antibiotic, bedaquiline exhibits significantly reduced efficacy against *M. abscessus* growing as a biofilm compared to planktonic growth, but that the efficiency of antibiotic penetration was not the main explanation for the different susceptibilities of MABC biofilms to treatment.

## Introduction

*Mycobacterium abscessus* complex (MABC) is a group of rapidly growing, nontuberculous mycobacteria (NTMs) frequently associated with pulmonary infections, but also skin and soft tissue infections (1, 2). MABC comprises of three subspecies: *M. abscessus* subsp. *abscessus*, *M. abscessus* subsp. *bolletii* and *M. abscessus* subsp. *massiliense* (3), which exhibit two colony morphotypes; rough and smooth, based on the absence or presence of the cell wall glycopeptidolipids (GPLs) (4). These species can form biofilms contributing to localised infections, especially in the lungs (4–6). Biofilms are defined as a community of aggregated bacteria enveloped in an extracellular matrix (ECM) (7–9). This growth mechanism renders them notoriously tolerant to antibiotics, contributing to the treatment recalcitrance of *M. abscessus*. This is often thought to be due poor penetration of antibiotic into the biofilm leading to subinhibitory concentrations reaching the bacteria.

Establishing standardized antibiotic susceptibility testing methods with good *in vitro*-*in vivo* correlation is a significant challenge in managing MABC infections as susceptibility testing outcomes do not align with treatment success against these infections (10, 11). In addition, the conventional minimum inhibitory concentration (MIC) testing, used to measure antibiotic efficacy against bacteria undergoing planktonic growth, does not align with the concentrations required to combat biofilms (12). Macrolides are crucial for treating MABC infections, but their effectiveness is limited due to macrolide resistant MABC strains (13–15). Alternative agents like Bedaquiline (BDQ) are being explored to address these challenges. BDQ inhibits the proton pump of ATP synthase, an enzyme that is required for cellular respiration (16, 17). This bromine-containing diarylquinoline shows promise, with low MIC values against MABC isolates offering a potential treatment option for these challenging infections (18–20)

In biofilms, the ECM provides stability to the biofilm structure but may also act as a protective layer and diffusion barrier inhibiting antibiotic penetration to access single cells. Advanced imaging techniques can play a valuable role in understanding the interaction and penetration of antibiotics into bacterial biofilms. For example, scanning electron microscopy (SEM) provides high-resolution imaging of the biofilm architecture at the level of individual bacterial cells (21) Studies using SEM have shown that biofilm cells exposed to antibiotics undergo morphological changes (22, 23).

Nanoscale secondary ion mass spectrometry (NanoSIMS) offers the ability to simultaneously detect up to seven isotopic masses at subcellular spatial resolution, enabling the mapping of labelled molecule distributions within cells. NanoSIMS is capable of distinguishing specific subcellular structures and has been combined with electron microscopy techniques to characterise cellular ultrastructure (24). This correlative approach has been used to study the activity of pyrazinamide (PZA) and BDQ on *Mycobacterium tuberculosis* within human macrophages, revealing how these drugs accumulate within bacterial cells and impact their efficacy (25) and to track the distribution of BDQ within individual cells in murine infected lungs revealing its localisation within intracellular bacteria and specific cell types (26)However, to date there have not been any studies investigating antibiotic penetration into NTM planktonic cells or biofilms using NanoSIMS.

This study aims to evaluate the efficacy of BDQ against MABC growing planktonically or as biofilms and assess its bactericidal action using a variety of readouts: minimal bactericidal concentration (MBC); minimal biofilm eradication concentration (MBEC); minimal biofilm inhibitory concentration (MBIC) and biofilm bactericidal concentration (BBC). These data are correlated with the results of the SEM and NanoSIMS imaging on the impact of BDQ on biofilm structure and its ability to penetrate biofilms and reach individual bacterial cells.

## Results

### MIC of BDQ against MABC (planktonic bacteria)

Drug susceptibility testing was performed using broth microdilution to determine the minimum inhibitory concentration (MIC) of BDQ for the MABC as planktonic bacterial cells. The species were exposed to a range of antibiotic concentrations, BDQ (0.0039-2 µg/ml) for an incubation period of 3 days. The MIC of BDQ for MABC was 0.25 µg/ml [Figure 1(A)], consistent with other reports (27, 28).

**Figure 1:**
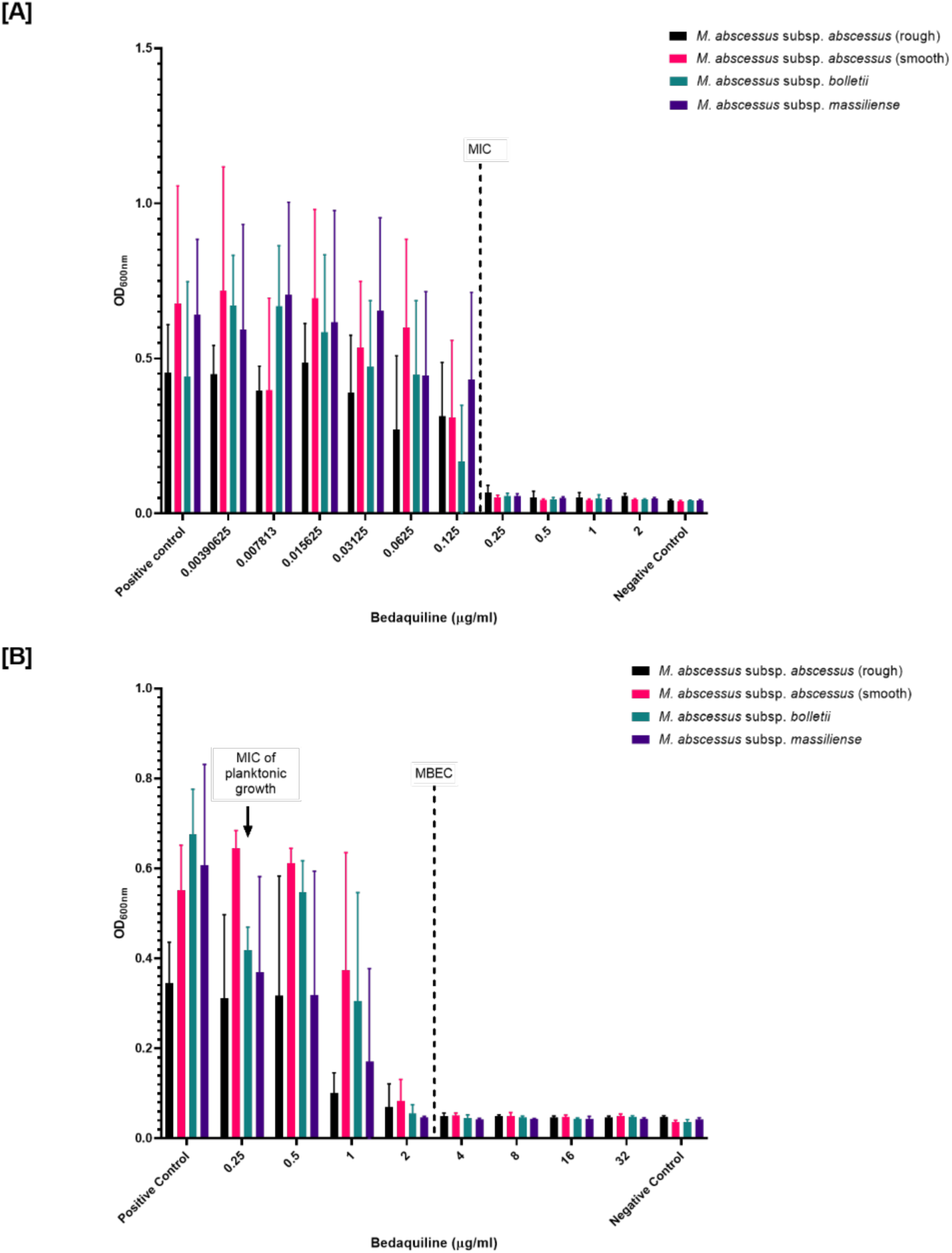
Determining the efficacy of Bedaquiline against MABC in planktonic growth and biofilms. **[A]** Antibiotic concentrations of BDQ were conducted in two-fold serial dilutions in a 96 well plate to determine the MIC. The OD_600nm_ in each well of the microtiter plate was measured after incubation at 37°C. The MIC was taken as the lowest concentration that inhibited MABC growth. The MIC for MABC was 0.25 µg/ml. **[B]** Using the MIC of the MABC as an initial concentration, the MBEC for BDQ against MABC biofilms were determined. Mature biofilms formed on the pegs of the device, were exposed to a two-fold serially diluted BDQ in a 96 well plate to determine the MBEC. After incubation for 24 hours at 37°C, the OD_600nm_ in each well of the microtiter plate was measured. The MBEC of MABC biofilms was 4 µg/ml. Results were expressed as the mean ± standard deviation (SD) of three independent experiments.

### Determining BDQ susceptibility against MABC biofilms

The efficacy BDQ to inhibit and/or eradicate MABC biofilms was tested. The mycobacterial species were exposed to 2-fold increasing antibiotic concentrations for 24 hours to determine the biofilm susceptibility endpoint parameters i.e. MBC, MBIC, MBEC and BBC, using the MIC value (0.25 µg/ml) for the planktonic cell as a starting concentration. The concentration of BDQ ranged between 0.25-32 µg/ml in the MBC, MBIC, MBEC and BBC assays.

Table 1 shows an overview of the MBC, MBIC, MBEC and BBC of BDQ determined for MABC biofilms.

**Table 1:**
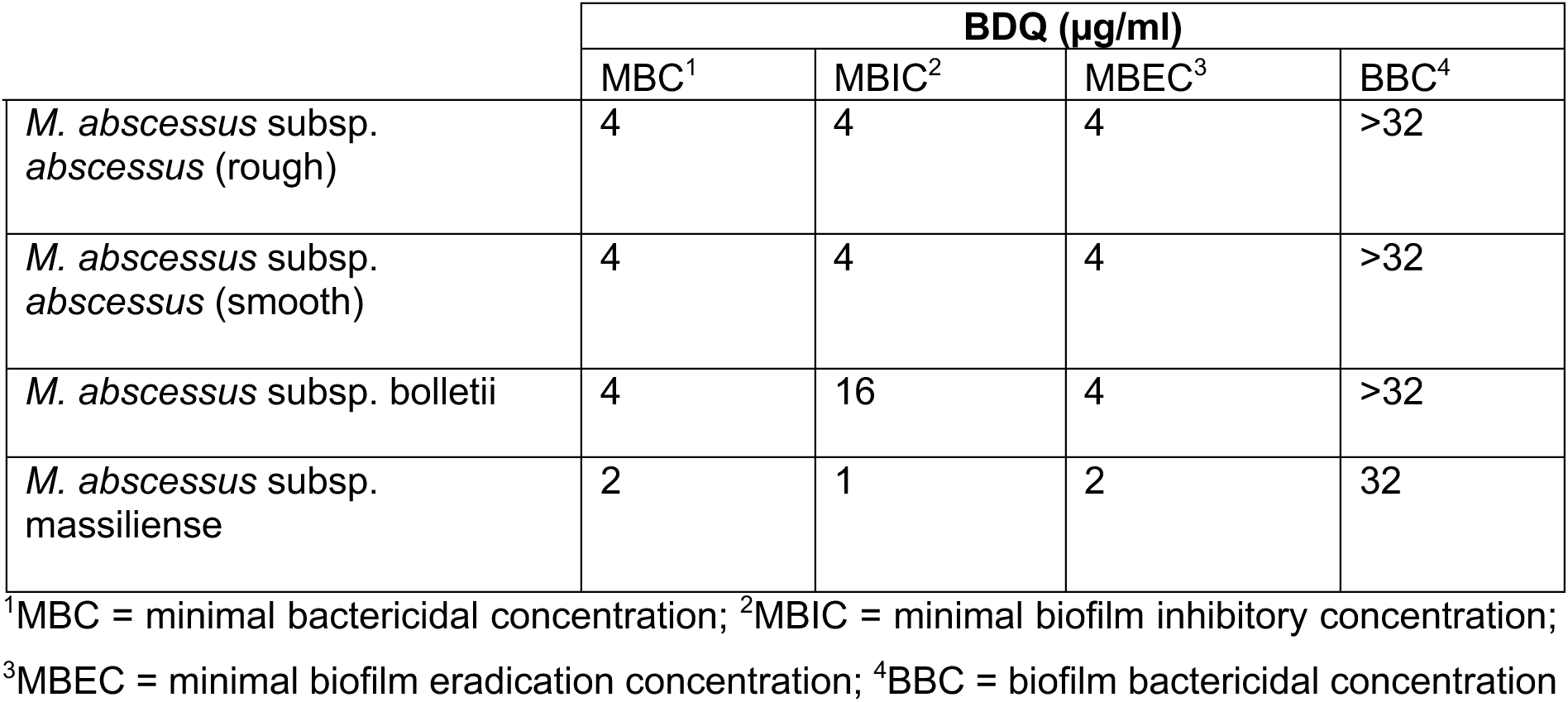
The MBC, MBIC, MBEC and BBC of BDQ against MABC biofilms using MBEC assay.

The MBC, MBEC, MBIC and BBC concentrations for MABC biofilms were significantly higher in comparison with the planktonic cells. The MBC for MABC was 4 µg/ml, except for *M. abscessus* subsp. *massiliense*, which was 2 µg/ml. MBIC values for *M. abscessus* subsp. *abscessus* biofilms (rough and smooth morphotypes) were consistent at 4 µg/ml, while *M. abscessus* subsp. *massiliense* biofilms had the lowest value (1 µg/ml). Conversely, *M. abscessus* subsp. *bolletii* biofilms required a higher inhibitory concentration (16 µg/ml). The MBEC for MABC (4 µg/ml) indicated BDQ had an inhibitory effect on biofilms [Figure 1(B)]. The BBC concentration (>32 µg/ml) across all *M. abscessus* subspecies biofilms revealed BDQ lacked a bactericidal effect (Supplementary Figures).

### Morphology and structure of MABC biofilms exposed to BDQ imaged by SEM

MABC biofilms were grown on sterile silicon chips and incubated for 7 days at 37°C. They were exposed to BDQ (4 µg/ml, based on MBEC results) for 24 hours before freeze-drying and SEM visualisation. SEM was used to assess the effect of BDQ on the morphology of the freeze-dried *M. abscessus* (subspecies) biofilms.

In untreated *M. abscessus* (rough) biofilms, SEM images revealed compactly embedded rod-shaped bacilli within a dense matrix [red arrows in Figure 2(Aii)]. However, with BDQ addition, SEM images showed less dense, porous *M. abscessus* (rough) biofilms compared to untreated samples [Figure 2(Aiv-vi)]. BDQ caused the breakdown of the dense matrix and altered the morphology of the bacilli from rod-shaped to round-shaped [indicated by a red circle in Figure 2(Avi)]. Similarly, *M. abscessus* (smooth) biofilms displayed intermingled bacilli within a highly dense, porous matrix [yellow arrow in Figure 2(Bi)]. The addition of BDQ to *M. abscessus* (smooth) biofilms showed a breakdown of the ECM and the release of individual bacilli [Figure 2(Biv-vi)]. BDQ also disrupted the cell membrane of *M. abscessus* (smooth), resulting in altered membrane structure [indicated by yellow arrows in Figure 2(Bvi)]. Untreated *M. bolletii* biofilms exhibited a network-like structure of bacilli within the matrix [Figure 2(Ci-iii)]. However, in treated *M. bolletii* biofilms, the network structure was disrupted with thread-like connections between bacilli and an increased number of elongated individual bacilli [green arrows in Figure 2(Cv)]. Like *M. abscessus* (rough), *M. massiliense* biofilms showed compactly embedded bacilli within a dense matrix [Figure 2(Di-iii)]. BDQ addition caused the ECM breakdown and the release of bacilli embedded within the matrix. In addition, BDQ caused destruction of the cell membrane in *M. massiliense*, resulting in alterations to its structure [indicated by blue arrows in Figure 2(Dvi)].

**Figure 2:**
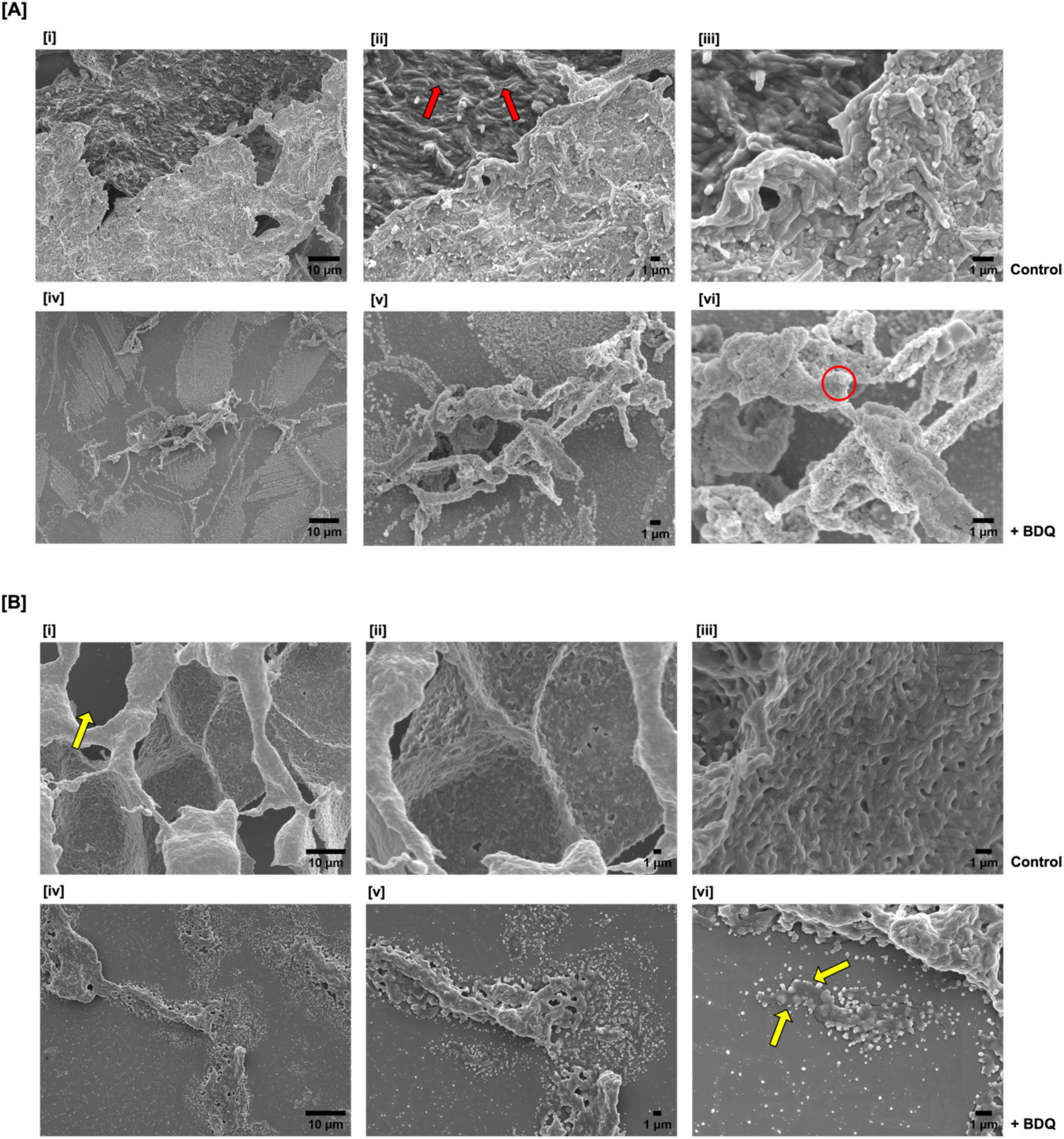
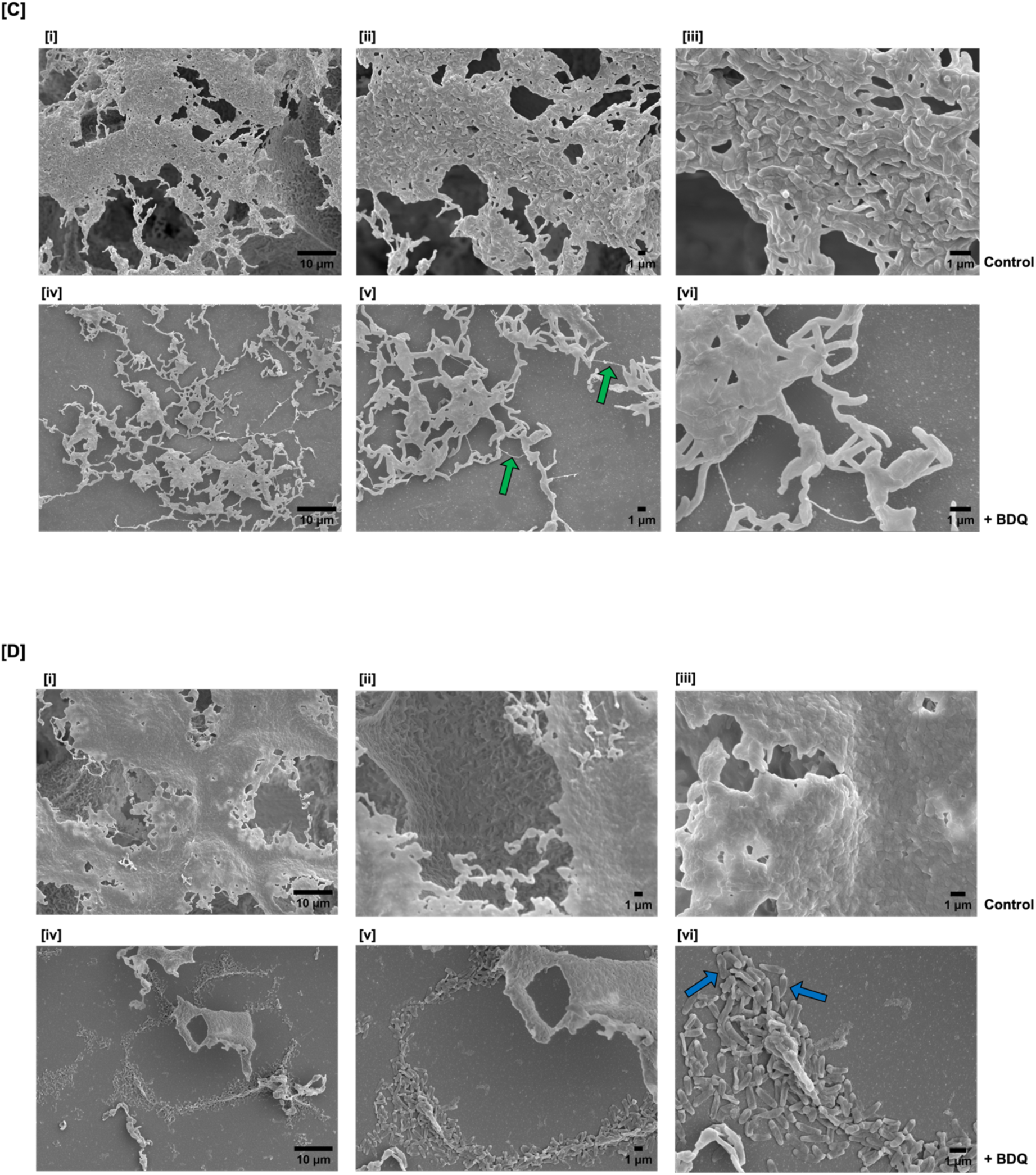
SEM analysis of MABC biofilms with and without BDQ. SEM micrographs illustrating the effects BDQ had on MABC biofilms. [(i-iii), increasing magnification] MABC biofilms without antibiotic showed bacilli embedded within an extracellular matrix (ECM). [(iv-vi), increasing magnification] After exposure to BDQ for 24 hours, MABC biofilms showed morphological changes within the structure. **[A]** *M. abscessus* subsp. *abscessus* (Rough) **[B]** *M. abscessus* subsp. *abscessus* (Smooth) **[C]** *M. abscessus* subsp. *bolletii* **[D]** *M. abscessus* subsp. *massiliense*.

### Distribution of Br^-^ at a single-cell level within MABC biofilms

The intensity of the total Br^-^ ion signal (a distinct marker for BDQ) was measured using NanoSIMS to determine the localisation of BDQ within the biofilms by summing the ^79^Br^-^ and ^81^Br^-^ ion images. The ^12^C_2_^-^ and ^12^C^14^N^-^ signals provided information on the structure of the cells within the MABC biofilms. Treated *M. abscessus* biofilms showed the Br^-^ ion signal within the cellular components, indicating the uptake of the lipophilic BDQ by cells of the biofilms, especially in the cell wall/matrix [Figure 3(A-D)].

**Figure 3:**
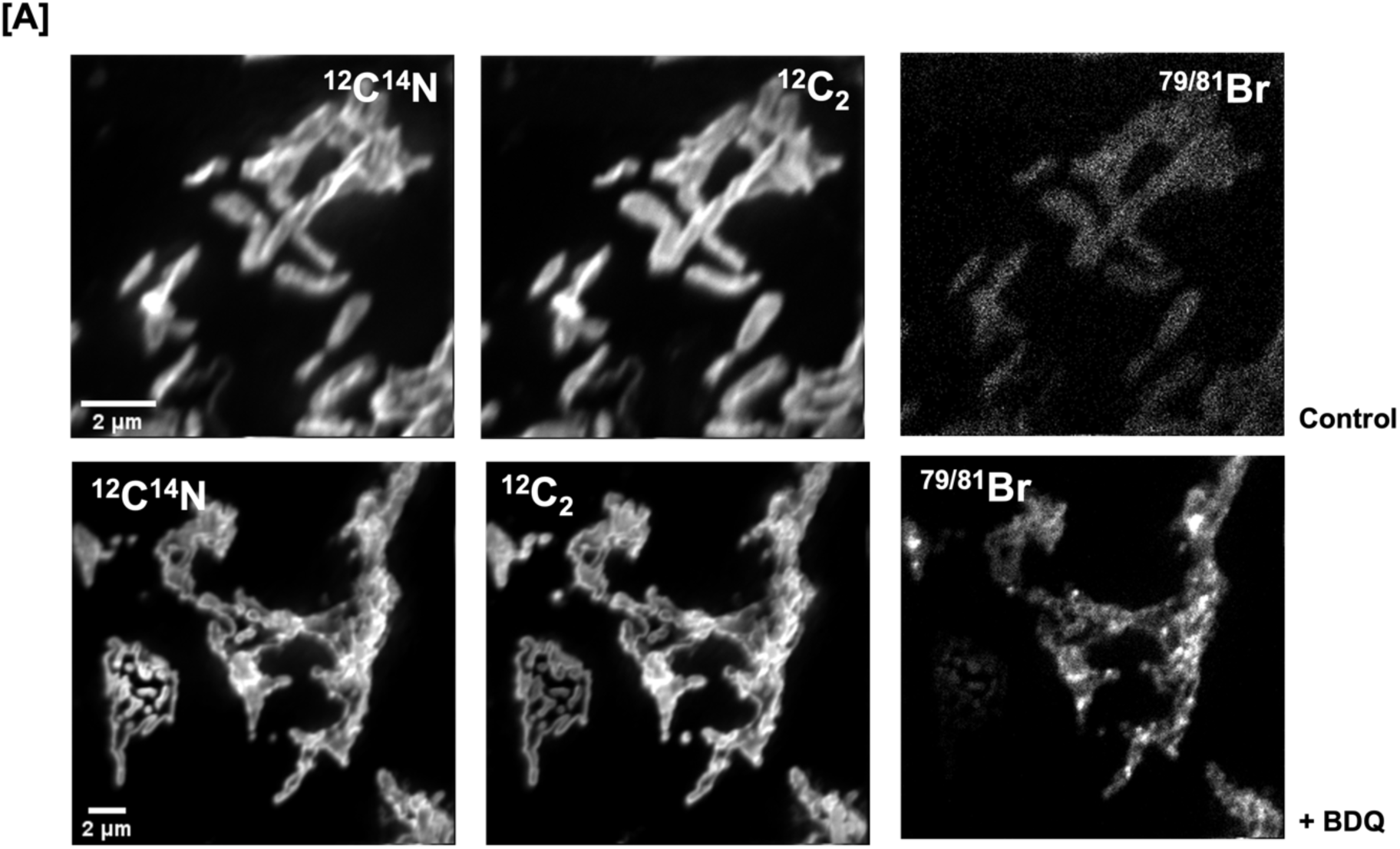
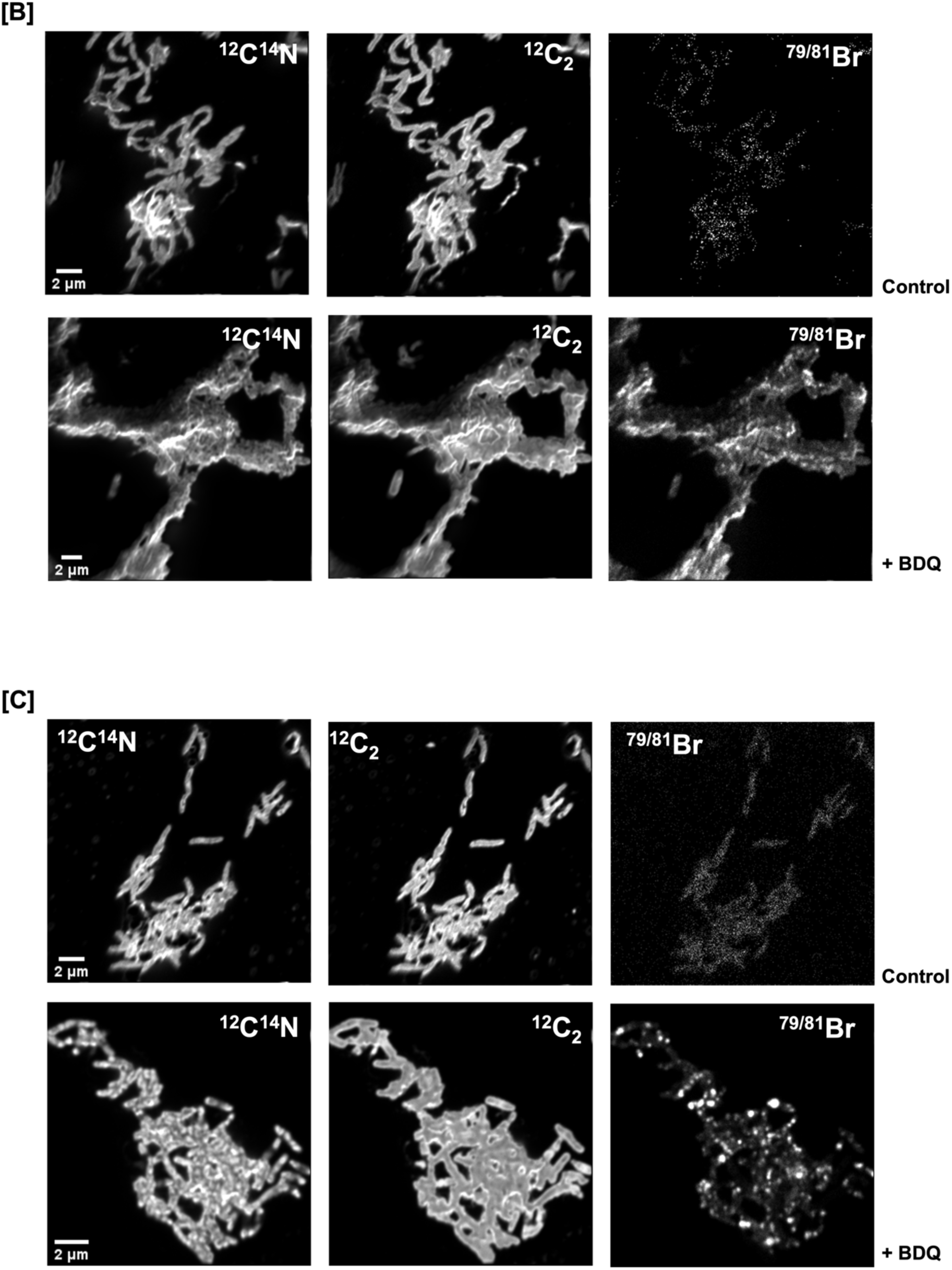
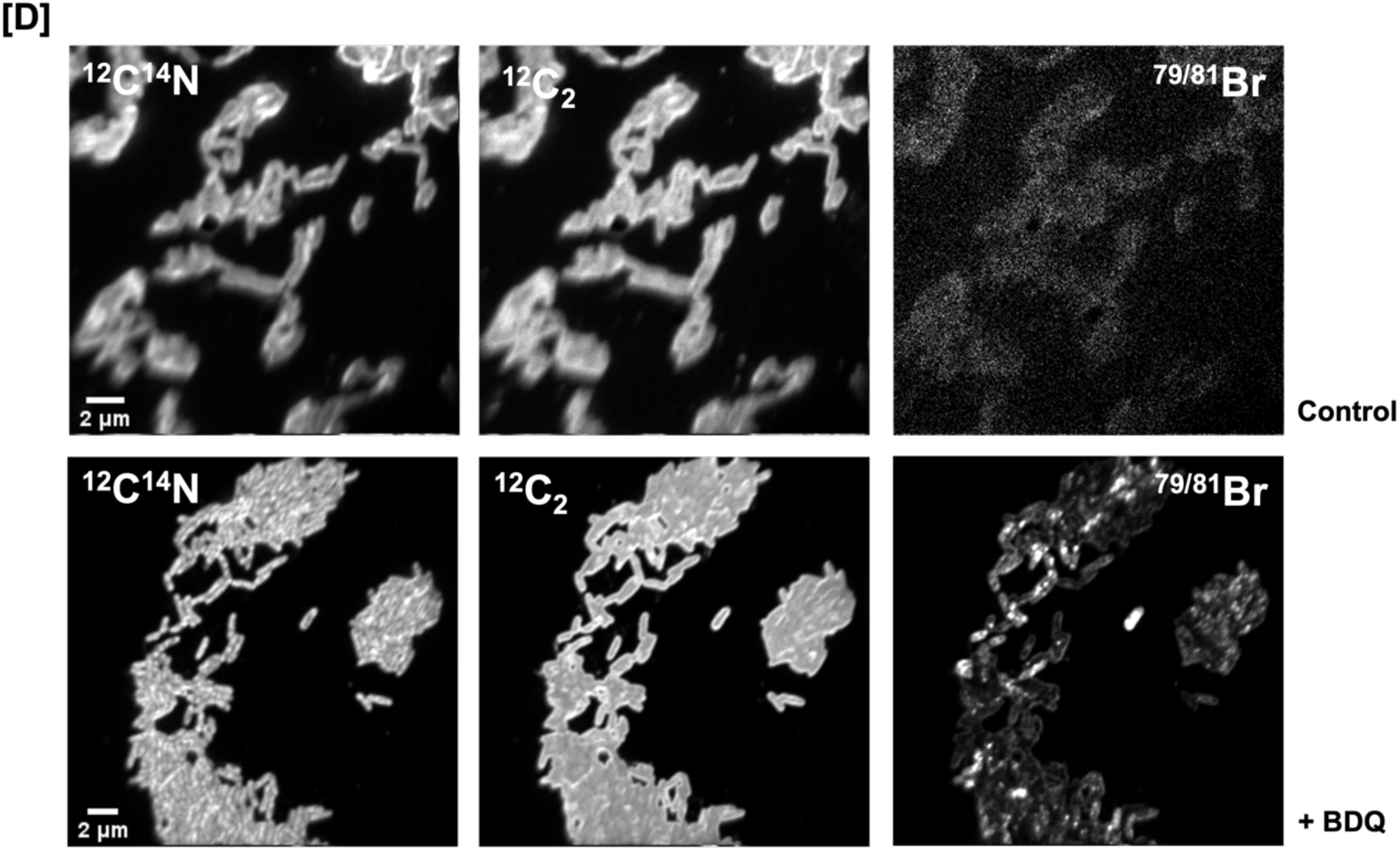
NanoSIMS micrographs of MABC biofilms treated with and without BDQ. Secondary ion images of cellular components of the biofilm (^12^C^14^N^-^ and ^12^C_2_^-^) and BDQ (Br^-^). Control MABC biofilms (Top Row) and MABC biofilms treated with BDQ for 24hrs (Bottom Row). **[A]** *M. abscessus* subsp. *abscessus* (Rough) **[B]** *M. abscessus* subsp. *abscessus* (Smooth) **[C]** *M. abscessus* subsp. *bolletii* **[D]** *M. abscessus* subsp. *massiliense*. Images are representative of randomly selected regions from three biological samples and three technical repeats.

### Quantitative analysis of the uptake of BDQ into MABC biofilms

Quantitative analysis was carried out using the ratio distribution Br^-^/^12^C^14^N^-^ and Br^-^/^12^C_2_^-^ to compare antibiotic penetration (based on the same BDQ concentration used) in the treated and untreated MABC biofilms. The quantitative analysis showed that there was significantly higher bromine uptake into the cell wall/matrix of *M. abscessus* subsp. *bolletii* biofilms compared with the other subspecies [Figure 4].

**Figure 4:**
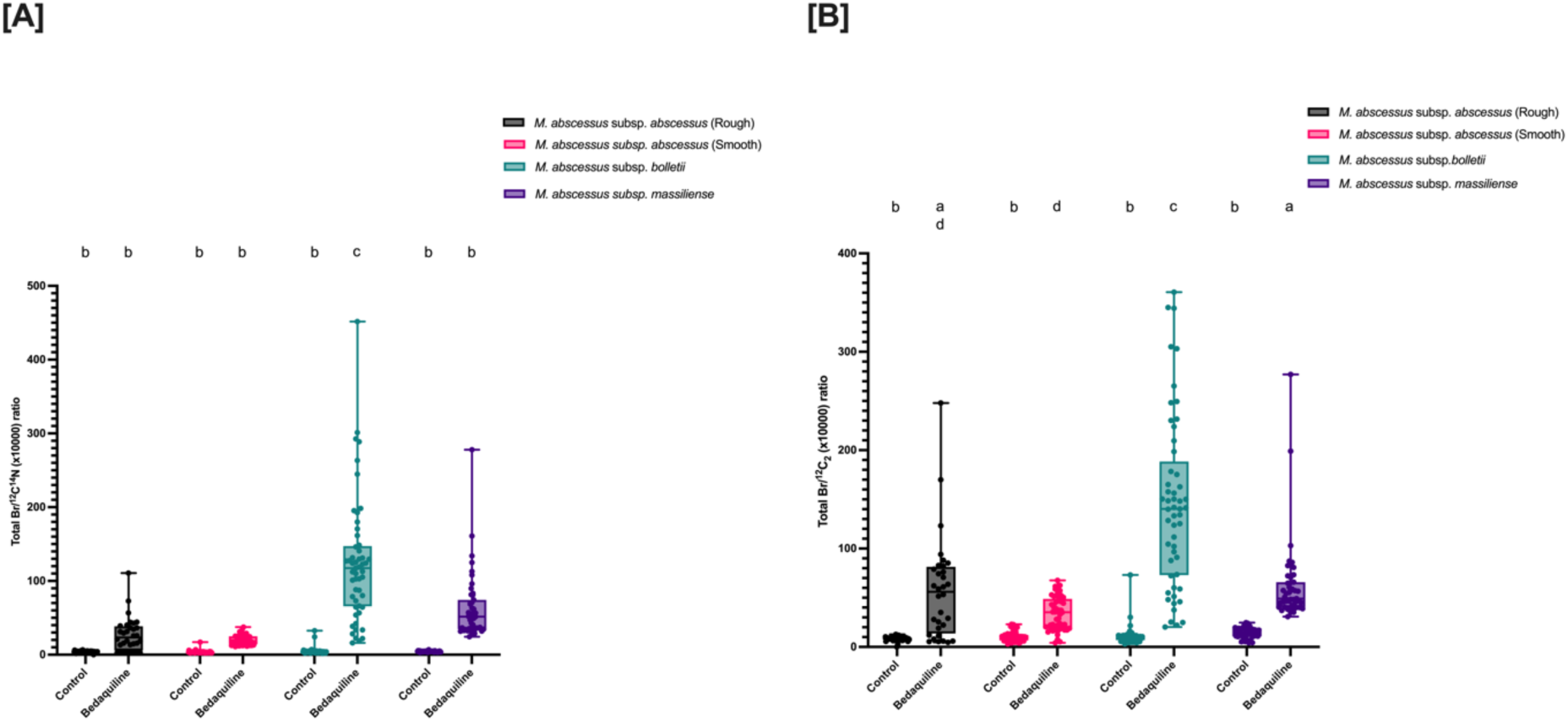
Quantitative analysis of BDQ distribution within the cellular components of MABC biofilms. Quantitative analysis using ratio distribution **[A]** Br^-^/^12^C^14^N^-^ **[B]** Br^-^/^12^C_2_^-^ of MABC biofilms with and without BDQ. Results were obtained from 32-61 individually segmented bacteria (from three biological samples and three technical repeats) and the *p*-values were calculated by using a Two-Way Anova. Pair-wise significant differences indicated by different letters.

## Discussion

Bedaquiline has been considered as a prospective antibiotic for the treatment of *M. abscessus* infections. Based on *in vitro* studies, BDQ has shown promising antibiotic efficacy against NTM species (29) and shown to be a good growth inhibitor at low doses against *M. abscessus* subspecies (19, 27, 30). However, it has been reported that BDQ does not have bactericidal activity against *M. abscessus in vivo* (19, 31, 32) which would be in-keeping with the findings in this study that *M. abscessus* subspecies exhibited reduced efficacy to BDQ as biofilm.

Antibiotic resistance is a hallmark of biofilm formation. Our results highlight that *M. abscessus* can form biofilms irrespective of morphotype. Based on the MIC of BDQ obtained for planktonic growth in this study, the MBEC assay was used to determine different biofilm susceptibility endpoint parameters (MBC, MBIC, MBEC and BBC) for subspecies of *M. abscessus*. These biofilm susceptibility endpoint parameters have been used as a guide for biofilm related infections. However, there are huge discrepancies between their definitions and parameters, therefore there is a need to have standardized definitions for these biofilm endpoints similar to the MIC (33). Based on previous studies (34–38)and for the context of this study, the MBEC was used as the biofilm susceptibility endpoint parameter to make comparison with the MIC. Our results showed that *M. abscessus* (subspecies) biofilms were inhibited at higher concentrations of BDQ compared with planktonic growth. In addition, BDQ had a bacteriostatic effect against the biofilms (2-4μg/ml) but needed a much higher concentration to have a bactericidal effect on the biofilms (≥32μg/ml). These findings suggest that while BDQ is considered a potential agent for *M. abscessus* infections due to its low MIC, it is essential to recognise the complexity introduced by biofilms given their genetic and phenotypic differences. Biofilms are much harder to eradicate compared with planktonic bacteria(39, 40).

*M. abscessus* subspecies exhibit either a rough or smooth morphotype based on the presence or absence of GPLs on the cell wall (41). Previous studies suggested that only the smooth morphotypes can form extensive biofilms whilst, the rough morphotypes produce either no or little biofilm but can still cause persistent infections (42–44). We studied the morphological heterogeneity of the MABC. SEM showed that both rough and smooth morphotypes were able to form extensive biofilms, contrary to prior findings. Consistent with other studies, *M. abscessus* (rough) also formed pellicles with a cording formation(34). It is believed that the MmpL protein is involved in the GPL biosynthetic pathway of the cell envelope, and that a mutation of *mmpL4a* gene causes a deficiency in GPL production and/or transport in the rough morphotype and the capacity to produce cords *in vitro* (45). The cording formation has been associated with increased pathogenicity and drug resistance (46). Studies have shown that cord-forming isolates such as *M. abscessus* (rough) are denser and contain a greater number of cells per unit compared with smooth isolates (47), which could reduce antibiotic penetration within the rough morphotype biofilm.

As the unique structural features of biofilms pose a barrier to antibiotic penetration contributing significantly to the emergence of functional antibiotic resistance, we used NanoSIMS imaging to examine the effects of BDQ on these biofilms. Cording formation of *M. abscessus* (rough) is thought to contribute to persistent lung infection and the recalcitrance of antibiotic treatment. Whilst this was consistent with high MBEC and BBC values observed for *M. abscessus* (rough), NanoSIMS analysis expressed as the Br^-^/CN^-^ ratio, did not indicate reduced uptake of BDQ into *M. abscessus* (rough) biofilms compared with the other *M. abscessus* subspecies.

The efficacy of antibiotic penetration into biofilms is not solely influenced by the cellular organisation or formation of the biofilm, but also by its cellular components and extracellular matrix (ECM) (48). The ECM contains lipids (GPLs, mycolic acids), polysaccharides, proteins and eDNA (49, 50). Generally, the rough morphotype has a low GPL expression, whereas the presence of the GPL in the smooth isolates has been linked with increased hydrophobicity and biofilm formation (51, 52). Based on the observation of the biofilm structure it can be deduced that *M. abscessus* subsp. *bolletii* likely has a smooth morphotype like *M. abscessus* subsp. *abscessus* (smooth) that expresses the GPL on its cell wall. NanoSIMS analysis showed an increased uptake of BDQ into *M. abscessus* subsp*. bolletii* biofilms (based on the Br^-^/CN^-^ ratio values) compared with the other *M. abscessus* subspecies. This is consistent with the observation that the network structure was disrupted in treated *M. bolletii* biofilms, and that the expression of GPL in *M. abscessus* subsp. *bolletii* may have caused the attachment of the lipophilic BDQ to the lipid-rich cell wall. There is very limited information regarding *M. bolletii* strains, specifically in relation to the rough and the smooth morphotypes and/or biofilm formation, despite these strains being associated with pulmonary infections (53–55). Studies suggest that GPL affects the mechanical properties of *M. abscessus* biofilms, whereby the smooth morphotype exhibited increased pliability compared with the rough morphotype (56). This characteristic could be a contributing factor in the increased uptake of BDQ into *M. bolletti* biofilms. However, this was not followed by increased efficacy of the drug; indeed, *M. bolletti* biofilms exhibited MBEC and BBC values no different to the other subspecies. Furthermore, as we found *M. abscessus* subsp. *abscessus* (smooth) biofilms exhibited least uptake of BDQ, there is a need to further explore the relationship between the structural properties (cords, aggregates) and the chemical composition of the biofilms (including the GPLs, mycolic acids) with antibiotic penetration.

Beyond the primary biofilm resistance mechanism of antibiotic penetration, another contributing factor may be the reduced uptake of antibiotics by oxygen-deprived bacteria within the biofilm. The biofilm architecture can create an oxygen and nutrient gradient. As a result, the outer layers of the biofilms are usually aerobic and metabolically active, whilst the inner layers become anaerobic and nutrient deficient. Most antibiotics are most effective against metabolically active bacteria, so the reduced growth rate on the inner layers of the biofilms could contribute to antibiotic resistance (42, 44, 46). SEM micrographs suggested that the presence of ECM hindered antibiotic penetration of BDQ to effective concentrations in the deeper biofilm layers. As a result, the outer layers of the biofilms exhibited morphological changes and in some of the mycobacterial species ECM degradation, while the deeper layers retained their structural integrity, showing little or no response to the antibiotic treatment.

Biofilm development can be a significant barrier in effective treatment of chronic NTM infections. This is also coupled with *M. abscessus* being intrinsically resistant and/or acquiring resistance to the currently available antibiotics. In this case, *M. abscessus* biofilms limited antibiotic penetration due to their colony morphotypes, biofilm architecture and the cellular composition of the biofilms. In addition, the slower growth rate, hydrophobic nature and/or the metabolic status of the mycobacteria within the biofilms (44, 57, 58), all likely contribute to increased antibiotic resistance. Due to the difficulties in the treatment of biofilm related *M. abscessus* infections, there is a need to know how to incorporate BDQ into these treatment regimens and to maximise its effect. It has been reported that BDQ was more effective in synergy with other antibiotics against disseminated NTM infection in patients co-infected with HIV (59). Further studies are needed to use combinational treatment therapy that include BDQ against *M. abscessus* biofilms and to determine their effects.

In conclusion, the combination of SEM and NanoSIMS imaging techniques allowed for the visualisation of the spatial arrangement of the biofilms, morphological effects of the antibiotic applied to the biofilms and tracking of antibiotic penetration at the single-cell level. These findings suggest that both the morphology and structure of the MABC biofilms and possibly the composition of the biofilm limited the efficacy of the antibiotics, but that antibiotic penetration alone could not account for the resistance of biofilm to treatment. These results point to the need for a deeper understanding of how the biofilm contributes to functional antibiotic resistance and that better therapeutic strategies need to do more than merely increase antibiotic penetration into biofilms.

## Materials and Methods

### Bacterial strains, media, and growth conditions

*M. abscessus* subsp. *abscessus* (*M. abscessus*)*, M. abscessus* subsp. *bolletii* (*M. bolletii*), and *M. abscessus* subsp. *massiliense* (*M. massiliense*) clinical isolates were a kind gift from the National *Mycobacterium* Reference Laboratory, Borstel, Germany. Liquid cultures of mycobacterial species were grown in Middlebrook 7H9 supplemented with 10% Albumin Dextrose Catalase (ADC) (Sigma-Aldrich) for MABC, 0.2% glycerol and 0.05% Tween 80 (Sigma-Aldrich) incubated at 37°C. Overnight cultures contained glass beads to mechanically prevent the clumping of mycobacterial isolates.

### Determination of Minimum inhibitory concentration (MIC) of planktonic MABC

Bedaquiline (BDQ) was purchased from Combi-Blocks (San Diego, USA) and was dissolved in dimethyl sulfoxide (DMSO) at 1mg/ml concentration and stored at -20°C. The minimum inhibition concentration (MIC) assay was performed using broth microdilution as outlined by the Clinical and Laboratory Standard Institute (CLSI) (CLSI, 2011) to determine the lowest antibiotic concentration that inhibited the growth of planktonic cells. Cation-adjusted Mueller-Hinton broth (CAMHB) with 5% OADC was used for the MIC assay of the MABC. The concentrations of BDQ ranged from 2 to 0.0039 µg/ml by serial dilutions in a 96-well microtiter plate. Bacterial cultures were adjusted to approximately 1x10^7^ CFU/ml for inoculation. A volume of 50 µl of the diluted inoculum was added to each of well and the final volume in each well was 100 µl. In the experiment, the negative control consisted of just media or antibiotic, whilst the positive control consisted of bacteria alone. The plates were sealed and incubated at 37°C for 3 days. Following incubation, the minimum inhibitory concentration (MIC) was determined by measuring the optical density (OD_600nm_) using a plate reader. The MIC is defined as the minimum antibiotic concentration to inhibit planktonic growth.

### Biofilm antimicrobial susceptibility of MABC

Biofilm antimicrobial susceptibility testing was performed using methods previously described (34–38) to determine the minimum biofilm eradication concentration (MBEC). The MBEC biofilm inoculator (Innovotech Inc., Edmonton, Canada) consists of a plastic lid with 96 pegs and a corresponding base. Bacterial cultures were adjusted to approximately 1x10^6^ CFU/ml using Sauton’s minimal media and 150µl was dispensed into each well of the MBEC inoculator. Biofilms were formed on the pegs after 3-5 days incubation for 37°C. The mature biofilms (peg lids) were transferred into a new 96 well plate containing serially diluted BDQ from the stock solutions (1mg/ml), which were incubated for 24hrs at 37°C. Following the antimicrobial challenge, the peg lid was rinsed twice in PBS and exposed biofilms were disrupted into recovery media (7H9-ADC-Tween 80-glycerol) by sonication for 10 mins. Negative controls contained just media/antibiotics and positive controls contained bacterial biofilms alone.

### Determination of the minimum bactericidal concentration (MBC) and minimum biofilm inhibitory concentration (MBIC)

To determine the minimum bactericidal concentration (MBC) against *M. abscessus* (subspecies) biofilms, 20 µl of broth suspension from each challenge plate well was combined with 180 µl of fresh CAMH with 5% OADC in a new 96-well plate, sealed, and incubated at 37°C for 3 days. The MBC, defined as the lowest antibiotic concentration that inhibited growth of dispersed cells from the biofilm compared with the growth control. This was determined by measuring optical density at OD_600nm_ using a plate reader.

Concurrently, in the challenge plate, biofilms released planktonic cells into antibiotic-containing broth during the challenge incubation period. Subsequently, a new non-pegged lid was placed on the 96-well challenge plate base and incubated at 37°C for 3 days. Following incubation, the minimum biofilm inhibitory concentration (MBIC) of BDQ was determined by reading the optical density of the challenge plate at OD_600nm_, where MBIC is the minimum antibiotic concentration inhibiting the growth of dispersed cells from the biofilm.

### Determination of minimum biofilm eradication concentration (MBEC) and bactericidal biofilm concentration (BBC)

Following the challenge plate, the MBEC peg lid was transferred to a plate containing 200 µl of neutralizer recovery media (CAMH with 5% OADC), then sonicated to dislodge biofilms from the pegs. Planktonic cells in the recovery media were serially diluted (10^0^-10^7^) in PBS with Tween 80, and 20 µl of each diluted sample were spotted on Middlebrook 7H11 agar. The plates were then incubated for 3-5 days at 37°C. To determine the BBC, the colonies were counted and expressed as Log_10_ CFU/ml. BBC is defined as the lowest antibiotic concentration killing 99.9% of biofilm – embedded bacteria compared to the growth control.

After the Log_10_ biofilm reduction, the volume of the neutralizer recovery media used for serial dilution was replaced with fresh media. The plate was covered with a new sterile non-pegged lid and incubated at 37°C for 3 days. After incubation, the MBEC was determined by measuring the optical density at OD_600nm_. MBEC is defined as the minimum antimicrobial concentration that eliminates the biofilm.

### Statistical analysis

All experimental assays were carried out at least three times and their averages were determined where necessary. All data acquired in the study were statistically analysed using GraphPad Prism version 9.1.1 for Windows, GraphPad Software, Boston, Massachusetts USA, www.graphpad.com, to determine the *p-* values and establish any correlation between untreated and treated MABC biofilms using two-way ANOVA. If within the data sets, the *p*-value is less than 0.05 it was considered statistically significant.

### Sample preparation of MABC biofilms for Scanning Electron microscopy (SEM) and Nanoscale secondary ion mass spectrometry (NanoSIMS)

*M. abscessus* and subspecies were grown in Sauton’s minimal media on sterile silicon wafer chips (Agar Scientific Ltd., Stansted, UK) within a 6-well plate. The plates were incubated at 37°C without shaking to allow biofilm formation. After the incubation period, BDQ was added to the mycobacterial biofilm samples based on the concentration obtained from the MBEC assay. PBS was added as a negative control to the samples. Both samples were incubated at 37°C for 24 hours. After the incubation with and without antibiotics, the mycobacterial biofilm samples were rinsed with ultrapure water. The samples were placed in a -80°C freezer overnight and then placed into the freeze dryer (Modulyo, Edwards, UK) at -40°C with a pressure of 60-80 mbar, overnight. Once the samples were dried, they were either sputter gold-coated at a thickness 9 nm using a Quorum (Laughton, East Sussex, UK) sputter coater for SEM or left to air dry for NanoSIMS analysis.

### Scanning Electron Microscopy (SEM)

The cell morphology of the freeze-dried mycobacterial biofilms on silicon chips before and after the addition of the antibiotic (24 hours) treatment was assessed by SEM. Conductive adhesive carbon tabs were added to aluminium specimen mount stubs (12.5 mm) (Agar Scientific) and the silicon chips with the biofilms were carefully attached onto the top. The biofilm samples were viewed with a field emission SEM (JSM-7100F, JEOL BV, Zaventem, Belgium) using the secondary electron detector with an accelerating voltage of 5kV and probe current of 6 pA. SEM experiments were carried out in triplicate for samples with/without antibiotics.

### Nanoscale Secondary Ion Mass Spectrometry (NanoSIMS)

Nanoscale Secondary Ion Mass Spectrometry (NanoSIMS) (CAMECA, Gennevilliers France) was used to map and detect ions that are specific to the antibiotic (Br^-^ – BDQ)] within the mycobacterial biofilms. In addition, NanoSIMS was used in detecting the elements that can be found within the mycobacterial species (^12^C^14^N^-^,^12^C_2_^-^).

The sample was bombarded by a primary ion beam from a Cs^+^ source at a potential of +8 kV. The sample is biased at – 8 kV to extract negative secondary ions. Thus, the net impact kinetic energy of the ion beam is 16 keV. The resulting secondary ions (^12^C^14^N^-^,^12^C_2_^-^,^79^Br^-^, ^81^Br^-^) are steered through an electrostatic sector where they are filtered for energy and then to a magnetic sector mass analyser and are separated by their mass-to charge ratio. The positions of a set of electron multiplier detectors, with single ion sensitivity are adjusted so that one detector records the signal from one selected secondary ion mass. Signal is acquired at each pixel while the primary Cs beam is scanned across the sample surface, forming images. NanoSIMS has a high spatial resolution of <50 nm which allows for the identification of isotopic composition within individual bacteria samples at subcellular resolution. Images were acquired at either 128 x 128 or 256 x 256 pixels with primary beam currents ranging from 0.5 – 1.0 pA.

### NanoSIMS Imaging analysis

The ion signals obtained from the images were quantified using ImageJ with the OpenMIMS plugin (github.com/BWHCNI/OpenMIMS, Harvard Center for NanoImaging). Image processing included image alignment, image summing, selecting the region of interest (ROIs) using the CN and/or C_2_ images, extracting ion count intensities from the ROIs, and statistical analysis (60) and adding the scale bars. For the visualization of the uptake of BDQ in the bacteria, a ratio metric image was produced with Br^-^/^12^C^14^N^-^ x 10000.

## Acknowledgements

This work was supported by the Engineering and Physical Sciences Research Council (EP/S51391X/1) and the UK Department of Science, Innovation and Technology and the National Measurement System program. We would like to thank Dr Florian Maurer from Diagnostic Mycobacteriology, National Reference Center for Mycobacteria for providing all of the Nontuberculous mycobacteria (NTMs) strains used. We thank David Jones, Sarah Barnett-Tucker and Andreas Lakovidis from MicroStructural Studies Unit (MSSU), the electron microscopy facility of the University of Surrey for all the support provided on the SEM. We thank Dr. G. Greenidge and Dr. G. Trindade for reviewing the manuscript.

## Author Contributions

All authors contributed to the conception and the design of this study. W.A. and G.M. contributed to the acquisition and analysis of data. All authors contributed to the writing of the manuscript.

## Competing Interest Statement

The authors declare no conflict of interest.

## Notes

### Competing Interest Statement

The authors have declared no competing interest.

