## Supplementary figures and images for "Variation in drug penetration does not account for the natural resistance of *Mycobacterium abscessus* biofilms to antibiotic"

### S1.tiff

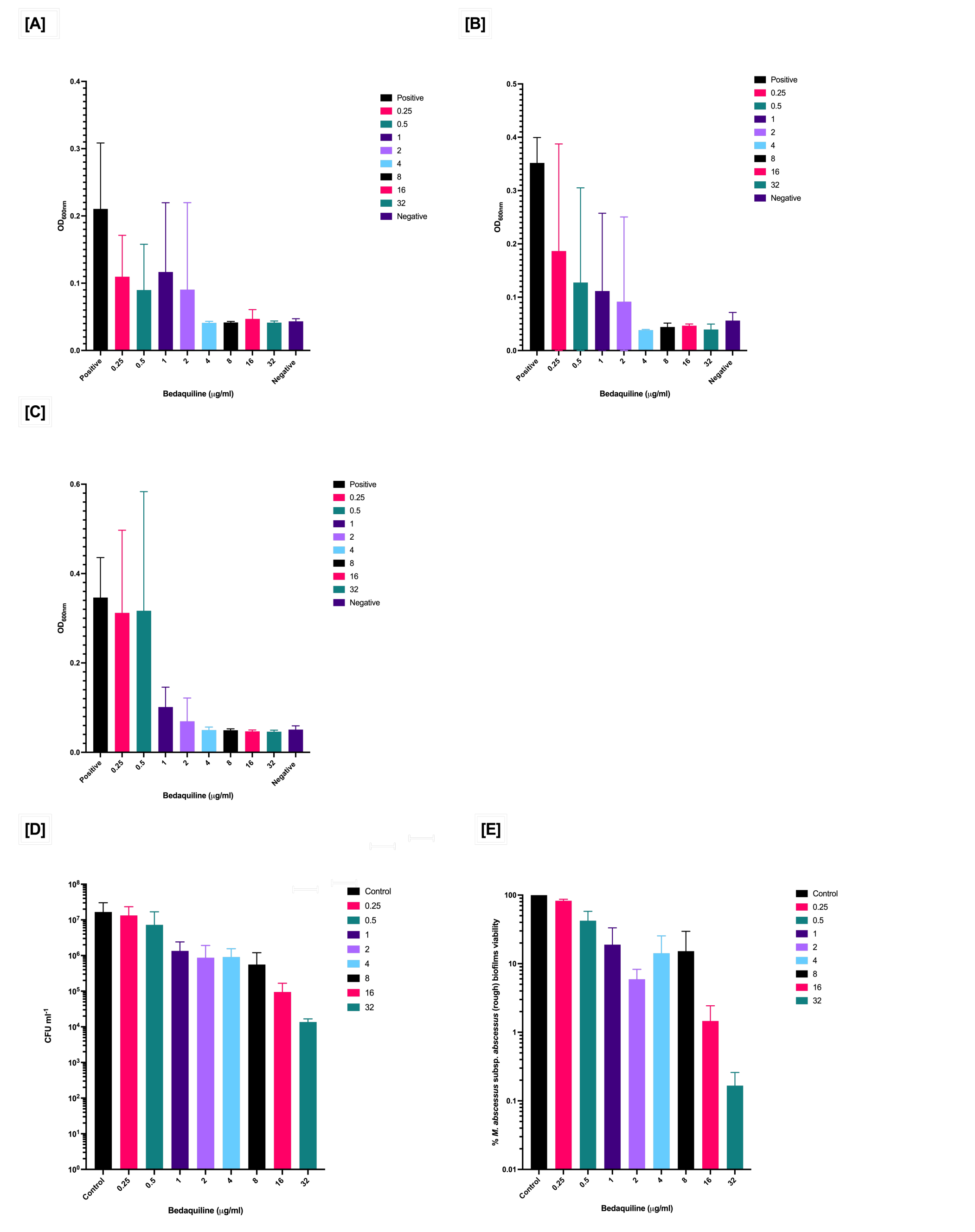

### S2.tiff

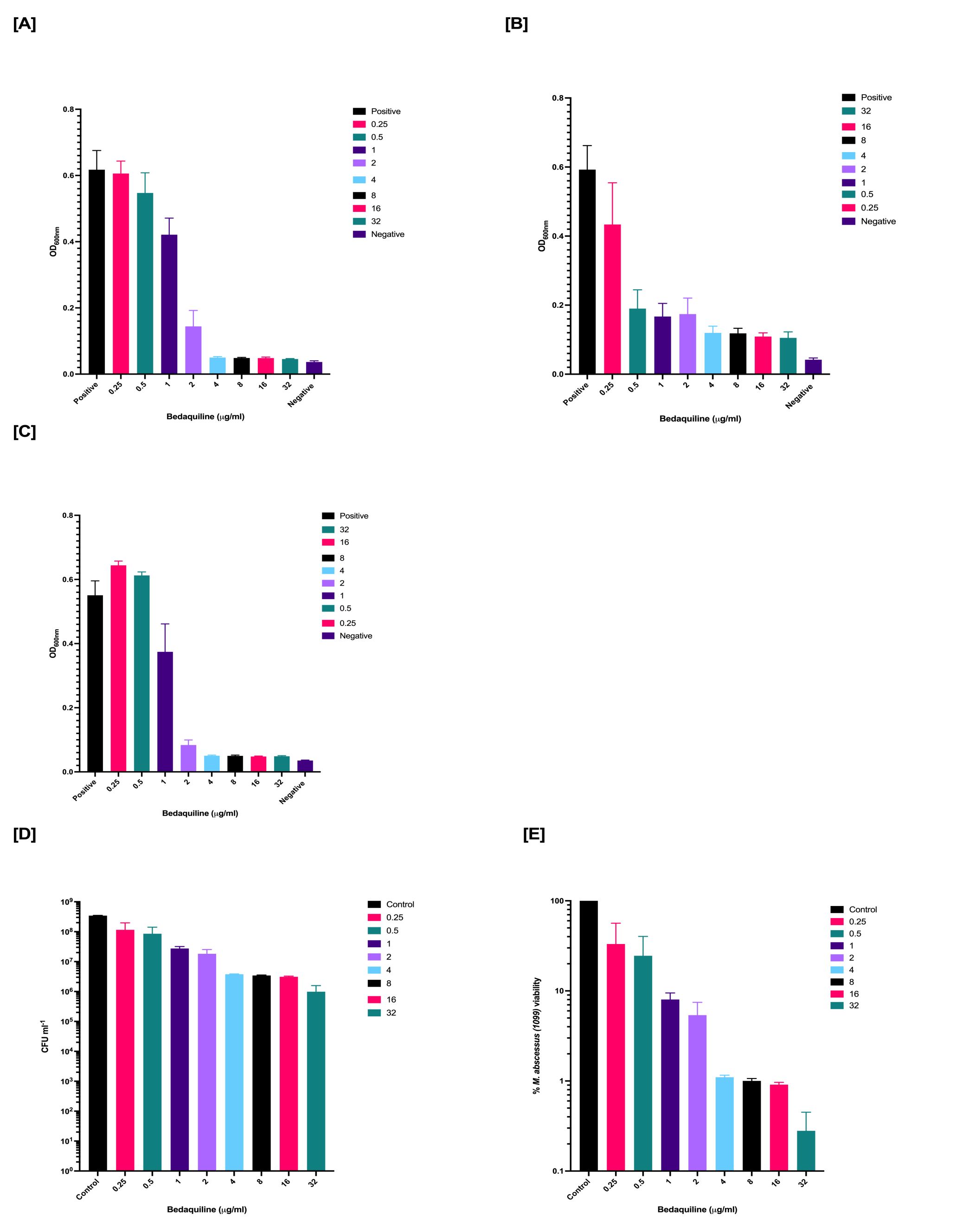

### S3.tiff

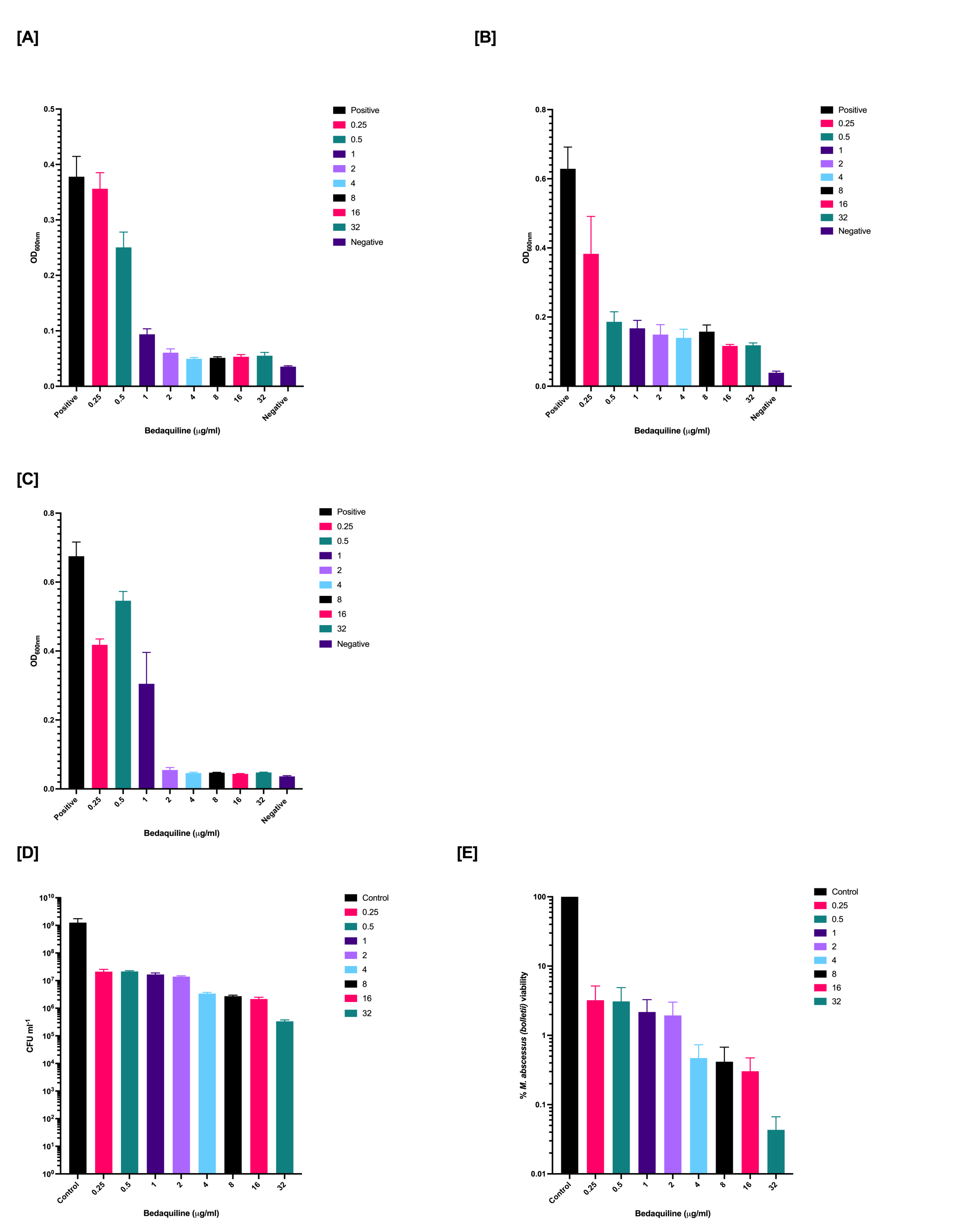

### S4.tiff

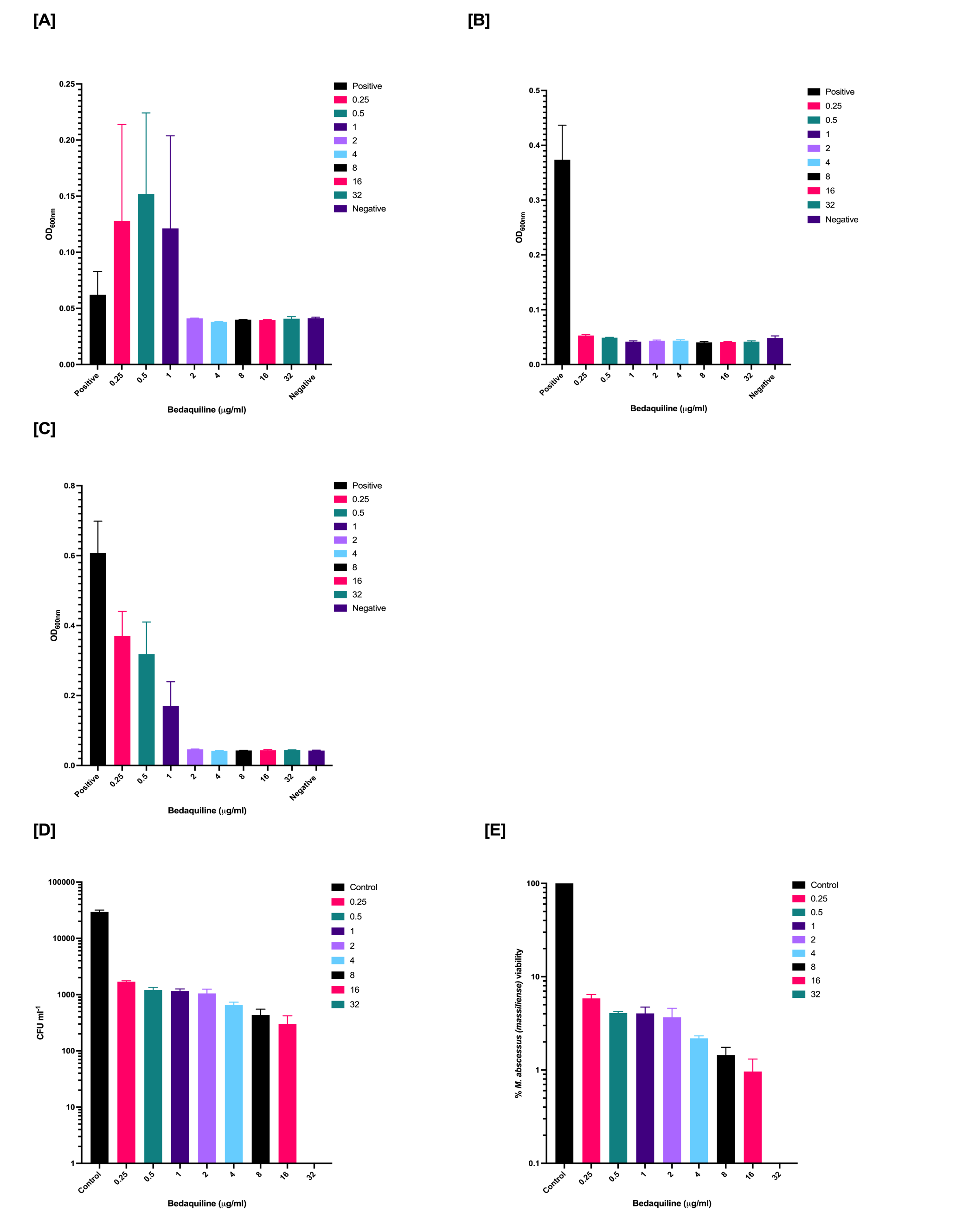
